# How life history can sway the fixation probability of mutants

**DOI:** 10.1101/050914

**Authors:** Xiang-Yi Li, Shun Kurokawa, Stefano Giaimo, Arne Traulsen

**Affiliations:** Department of Evolutionary Theory, Max Planck Institute for Evolutionary Biology, August-Thienemann-Straße 2, 24306 Plön, Germany; Division of Natural Resource Economics, Graduate School of Agriculture, Kyoto University, Oiwake-cho, Kitashirakawa, Sakyo-ku, Kyoto 606-8502, Japan

**Keywords:** Demography, Evolutionary dynamics, Population structure, Reproduction, Survival

## Abstract

In this work, we study the effects of demographic structure on evolutionary dynamics, when selection acts on reproduction, survival, or both. In contrast with the previously discovered pattern that the fixation probability of a neutral mutant decreases while population becomes younger, we show that a mutant with constant selective advantage may have a maximum or a minimum of the fixation probability in populations with an intermediate fraction of young individuals. This highlights the importance of life history and demographic structure in studying evolutionary dynamics. We also illustrate the fundamental differences between selection on reproduction and on survival when age structure is present. In addition, we evaluate the relative importance of size and structure of the population in determining the fixation probability of the mutant. Our work lays the foundation for studying also density and frequency dependent effects in populations when demographic structures cannot be neglected.

## 1 Introduction

The emergence and subsequent dynamics of mutants play important roles in determining the trajectory of evolution (Crow and Kimura, 1970). The fate of a mutant is strongly influenced by its relative fitness. The fitness of a mutant, or rather, the fitness landscape of the whole population is not static, but ever changing under influences of many factors, including density and frequency dependent effects (Nowak and Sigmund, 2004).

Depicting the trajectories of survival and reproduction along the lifespan, life history is well known to have great influence on evolution in general (Vindenes et al., 2009; Charlesworth, 2001; Nunney, 1991, 1996). It has been recognised being of fundamental importance in determining the trajectory of evolutionary dynamics (Wahl and DeHaan, 2004; Lambert, 2006; Hubbarde et al., 2007; Parsons and Quince, 2007a,b; Wahl and Dai Zhu, 2015). In addition, life history also modulates the actual effect that an increase in survival or reproduction has on the growth rate of the population, and it accounts for part of the effective size of the population (Charlesworth, 1994; Engen et al., 2005). For the frequency dependent case, it has been shown that life stage dependent strategic interactions can promote diversity and push strategic behaviours away from the equilibrium determined by game theoretic interactions alone (Li et al., 2015a). However, it is not obvious if certain demographic structures of the population would help or hinder the spread and fixation of a beneficial mutant. This is the problem of our interest in the present work, which may be the first step towards further studies of stochastic effects of the frequency dependent case.

Pioneer work on this topic includes seminal papers by Felsenstein (1971) and Emigh (1979a,b). One major goal of this previous work was to understand the evolutionary dynamics in populations with overlapping generations. Overlapping generation models are biologically more relevant to many species of interest including humans, but they are inevitably associated with increased complexity, due to age or stage structure of the population. To fully capture the demographic architecture of age-structured populations, not only the absolute number and frequency of the mutant matter, but also its distribution across different age classes. One important concept to note here is the reproductive value. The reproductive value of an individual was originally defined by Fisher (1930) as “to what extent will persons of this age (or sex), on average, contribute to the ancestry of future generations”. It does so by accounting for the remaining number of offspring an individual will produce, discounted by the increase of population size at the time of reproduction of this offspring. Fisher discovered that in a linear model, the total reproductive value in a population grows exactly exponentially regardless of whether the population age distribution has reached the demographically stable state. Thereafter, in population genetics models of age structured populations, it is of great importance to look at the dynamics of reproductive value weighted frequencies of alleles (Crow, 1979; Engen et al., 2005, 2009b; Vindenes et al., 2009). This reduces the typically highly dimensional problem of dealing with alleles flowing through multiple age classes to the lesser dimensional problem of tracking allele frequencies with the appropriate weights and essentially neglecting explicit age structure. More comprehensive explanations and examples are presented in the book of Caswell (2001). In populations with demographic structure, individuals of different age have different reproductive value. The evolutionary fate can be summarized by the fixation probability of a mutant, i.e. the probability that it ultimately takes over the entire population. This fixation probability can vary greatly in populations with overlapping generations with the fraction of young individuals in the population.

The pioneer researchers have made great analytical contributions under various simplifying assumptions, such as large population size (Felsenstein, 1971; Emigh, 1979a,b), neutral (Emigh, 1979a) or weak selection (Emigh, 1979b), and extreme demographic structures (Felsenstein, 1971; Emigh, 1979a,b), etc. Under these conditions, good approximations can be made to facilitate mathematical analysis. For example, if the sizes of subsequent age classes differ greatly or are very close to each other, it is cogent to approximate hypergeometric sampling with binomial sampling in order to capture the process of individuals entering the next age class. With impressive analytical dexterity, the pioneers successfully summarized the vast complexity into a few easily understood parameters, such as the “effective population number” (Felsenstein, 1971) and the “average reproductive value” (Emigh, 1979a). They used these parameters to describe and analyze evolutionary dynamics.

Despite the beautiful analysis, the original work did not achieve the impact it deserves in our opinion. This is probably due to the inevitable mathematical complexity or somewhat limited applications, dictated by the stringent underlying assumptions. With the help of current increased computational power, we show that there are still surprises left uncovered in the original model first described in Felsenstein (1971), even in the simplest case with only two age classes in a well-mixed population under constant selection.

In the following, we first describe the model and recall the fixation probability of a neutral mutant in the first age class as a benchmark. Then we analyze the pattern of fixation probability of a beneficial mutant in populations with diverse demographic structures. We explore one by one the evolutionary consequences of selecting on reproduction, survival and both. Further, we evaluate the relative importance of population structure and size in determining the fixation probability of a beneficial mutant. Finally, we show that unexpected non-monotonic patterns of fixation probability hold in populations with multiple age classes.

## 2 Model description

### 2.1 Life history and population updating rules

First, we describe the life history features of individuals and population updating rules, as illustrated in Figure 1. In contrast to the original model of Felsenstein (1971), we start from analyzing the simple case of only two age classes (and then go beyond it in section 7). In addition, we include different targets of natural selection, such as selecting on reproduction, survival or both. Consider a haploid population in which individuals can live up to age two at maximum. The numbers of individuals in age-one (young) and age-two (old) are constant, denoted as *N*_1_ and *N*_2_, respectively. In each time step, all individuals produce large amounts of offspring, proportional to their fitness. Among them, only *N*_1_ survive and become the next generation of young individuals. Similarly, only *N*_2_ of the young individuals at the previous time step survive to enter the old age class. All old individuals die and are removed from the population. The recruitment of young individuals from the pool of new born offspring is treated as a process of sampling *with* replacement (similar to the classic Wright-Fisher process), while the recruitment of old individuals from young ones is treated as sampling *without* replacement. The fitness of the mutant is assumed to be constant and greater than the fitness of the wild type, which is normalized to one. Selection may act on reproduction (fitter individuals have more offspring), survival (fitter individuals have higher probability to survive to the next age class), or a combination of both.

**Figure 1:**
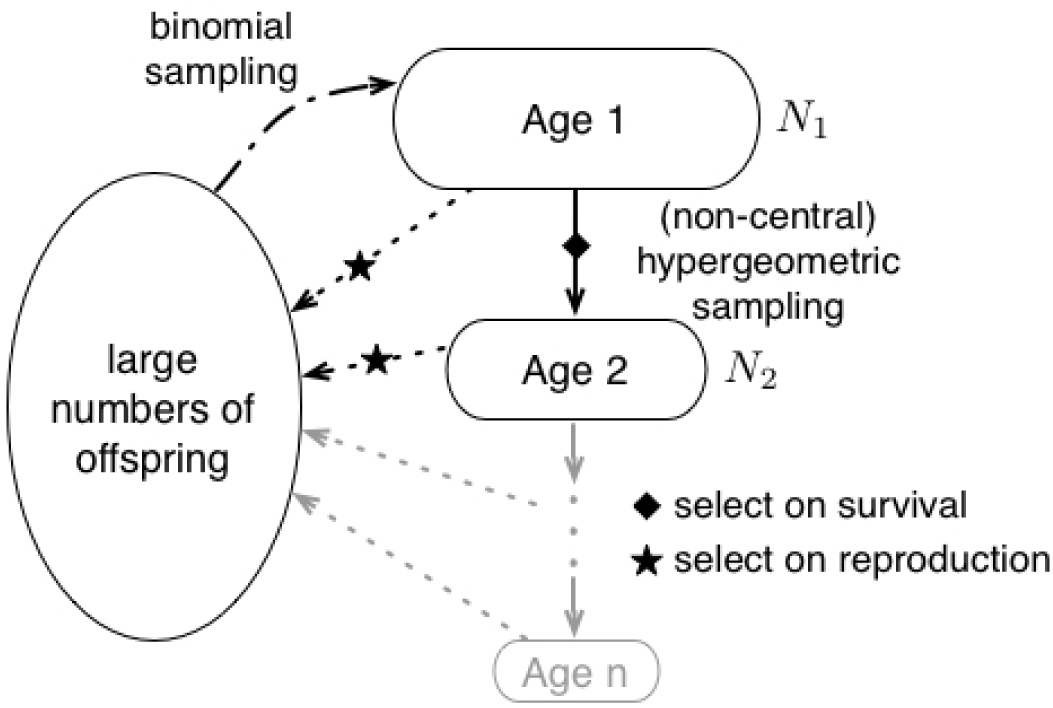
Population updating rules. The number of individuals in the first and second age classes are fixed (*N*_1_ and *N*_2_). Individuals produce large numbers of offspring proportional to their fitness, if selection acts on reproduction. Otherwise they all produce the same numbers of offspring. Offspring enters the first age class following binomial sampling, assuming that the number of offspring is much greater than *N*_1_. Entering of the second age class of the age-one individuals follows hypergeometric sampling. If selection acts on survival, fitter individuals have a higher probability to survive to the next age class. Consequently, the sampling process becomes non-central hypergeometric sampling. Compared to the original model described by Felsenstein (1971), we use only two age classes for simplicity, while the original model has an arbitrary number of age classes *ω* (we discuss this case in section 7). On the other hand, we explore the difference of selection on reproduction, selection on survival, and selection on both. These different sources of selective forces have not been considered in Felsenstein’s original model.

### 2.2 Fixation probability of a selectively neutral mutant

We denote the state of the population at time *t* as a tuple 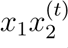, in which *x*_1_ is the number of mutants in the young age class, and *x*_2_ is the number of mutants in the old age class. The transition probability from 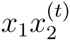 to 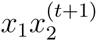 is

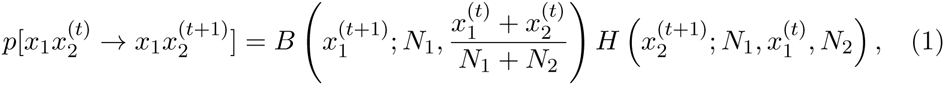
where 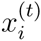 is the number of mutants in age class *i* at time *t*, *B*(*k; n, q*) = 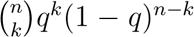 denotes the probability of obtaining *k* mutants in *n* draws with replacement, and *q* is the frequency of mutants in the population. We use this binomial sampling to model the process of picking *N*_1_ individuals from the offspring pool in order to form the new young age class. Similarly, 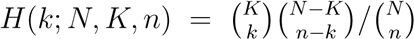 denotes the probability of obtaining *k* mutants in *n* draws without replacement, in which *N* is the total number of individuals, and *K* is the current number of mutants. We use this hypergeometric sampling for drawing *N*_2_ young individuals that enter the old age class in the next time step. There are (*N*_1_ + 1)(*N*_2_ + 1) different states of the age-structured population, therefore the state transition matrix 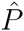 (with elements 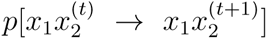) has the dimension (*N*_1_ + 1)(*N*_2_ + 1) × (*N*_1_ + 1)(*N*_2_ + 1). The fixation probability vector 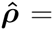 (*ρ*_00_, *ρ*_01_,…, *ρ*_0*N*_2__, *ρ*_10_, *ρ*_11_,…, *ρ*_1*N*_2__,…, *ρ*_*N*_1_0_, *ρ*_*N*_1_1_,…, *ρ*_*N*_1_*N*_2__) contains the fixation probabilities of the mutant type from each of the (*N*_1_ + 1)(*N*_2_ + 1) distinct population states.

The transition matrix 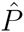 is stochastic and has unique eigenvalue 1. Due to the absence of mutation, we have *ρ*_00_ = 0 and *ρ*_*N*_1_*N*_2__ = 1. In practice, we can thus remove the first row and first column of the transition matrix 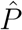 to form a new transition matrix *P* which corresponds to the fixation probability vector ***ρ***, which is 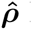 having the first element 0 removed. To calculate ***ρ***, we compute the eigenvector that corresponds to eigenvalue 1 of the transition matrix *P* numerically, and then normalize it by setting *ρ*_*N*_1_*N*_2__ to 1.

As a benchmark for later comparisons, we show in Figure 2 the fixation probability of a single neutral mutant arising from the young age class. We can see that a neutral mutant is more likely to reach fixation in populations with more old individuals than in populations with more young individuals.

**Figure 2:**
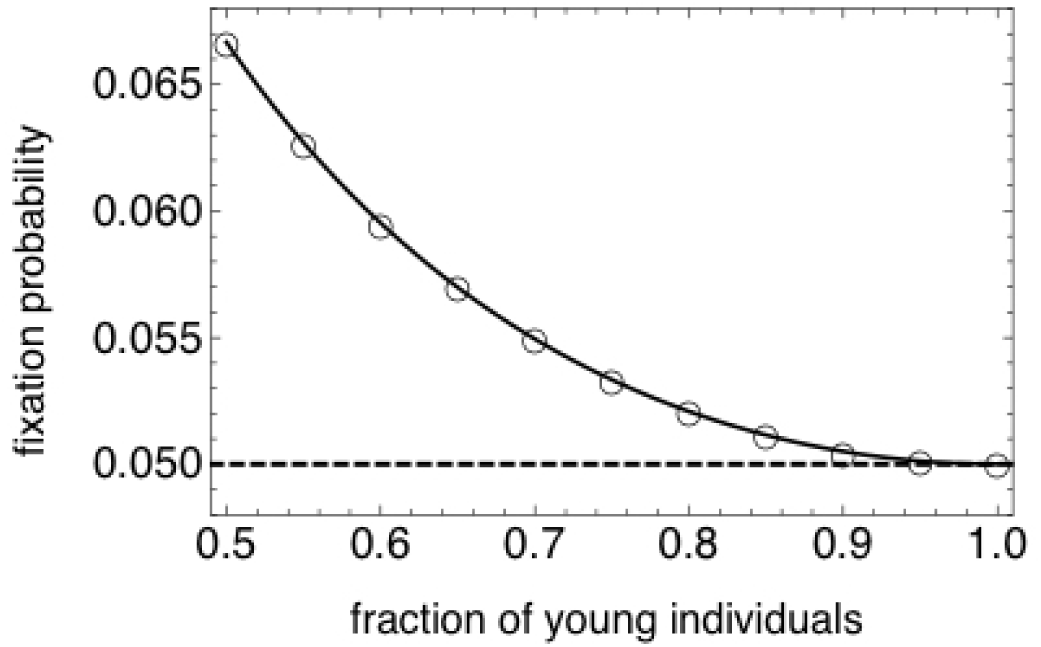
Fixation probability of a neutral mutant in a population of total size 20. Symbols are results from numerical calculation (based on the transition matrix), the solid line is the known analytical solution (see Appendix A). The dashed line marks the fixation probability when age structure is absent. There are two age classes in this example, therefore the fraction of young individuals ranges from 0.5 to 1. The neutral mutant has higher probability to reach fixation in populations with more old individuals than in populations with more young individuals. A Mathematica notebook file for generating this figure can be found in the Supplementary Information.

A natural explanation of this pattern is the differentiated reproductive values in the population. In an age-structured population, the fixation probability of a neutral mutation equals the initial frequency of the mutant sub-population weighted by reproductive values (Emigh, 1979a). The reproductive value of an individual of a given age expresses the contribution of this individual to the future ancestry of the population. We can use a matrix population projection model to compute the reproductive values of different ages (Caswell, 2001). This model is a square matrix that describes the vital rates (i.e. fertility and survival) of the population organized by demographic class, see Eq. 2 for example.

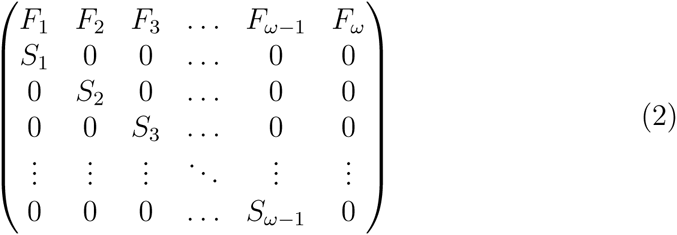

In the case of an age-structured population, the first row of the projection matrix contains the fertilities of each age class. The fertility *F*_*k*_ at the top of the *k*-th column gives the number of offspring born to an individual in age class *k* which successfully enters the first age class at the next time step. The subdiagonal of the matrix contains the age-specific survival probabilities. *S*_*k*_ stands for the fraction of individuals in age class *k* that will enter age class *k* + 1 at the next time step. All other matrix entries are zero. The matrix population model can be right-multiplied by the population state vector. The result of this multiplication is the projection of the population state to the next time step. If this operation is iterated, the vector representing the population state eventually becomes proportional to the leading right eigenvector of the matrix population model. At this point, the population is in a demographically stable state and its growth rate corresponds to the leading eigenvalue of the matrix. With appropriate scaling, the leading left eigenvector gives the reproductive value of each age class (Caswell, 2001). As this model only involves matrix algebra, it is entirely deterministic. In our case, the matrix population model is a 2 × 2 matrix, since we have only two age classes. As we assume a constant size, the leading eigenvalue of this matrix must be one.

To illustrate our approach, let us first consider the case of a neutral mutant, in which mutant and wild type have the same fitness. Using the description of the life history in our population given above, our matrix population model is written as:

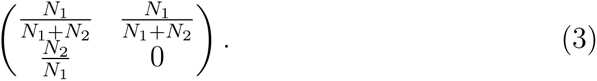

The right eigenvector of this matrix corresponding to the eigenvalue of one is 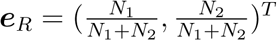. This vector is already scaled to give the stable age distribution in the asymptotic population. The corresponding left eigenvector 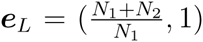 gives the reproductive values. With these two vectors, we can now give reproductive value weights to the initial frequency of a neutral mutant of young age in our population at demographic stability. The mutant’s reproductive value thus equals to 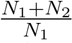. The reproductive value in the total population is the population size *N* multiplied by the scalar product *e*_*L*_ · *e*_*R*_, which is the total reproductive value at stability in the unit size population, *N*(*e*_*L*_ · *e*_*R*_) = *N*_1_ + 2*N*_2_ (Caswell, 2001). Taking the ratio of the initial mutant reproductive value to the total reproductive value in the population, we obtain 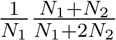, which corresponds to the fixation probability of the mutant (Emigh, 1979a). Observing that *N* = *N*_1_ + *N*_2_ is fixed, the derivative of this last quantity with respect to *N*_1_ is 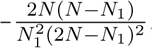, which is strictly negative. Therefore, the fixation probability of a neutral mutant decreases as the fraction of young individuals increases.

Another way to understand this pattern is to calculate the age-structure dependent fixation probability directly (Charlesworth, 1994). With the fraction of young individuals in the population *ƒ* = *N*_1_/*N*, the fixation probability of a single mutant in a population of size *N* is (*Nf*(2 − *ƒ*))^−1^, which is identical to our result above. A derivation of the fixation probability with this approach can be found in Appendix A.

To set the basis for later comparisons, we have first focused on the fixation probability of a neutral mutant in the first age class. In the following, we focus on the population dynamics and fixation probability of a beneficial mutant that has a constant selective advantage *r* > 1. This corresponds to the traditional notation of *r* = 1 + *s* in the classic population genetics literature. We explore one by one the effects of selection on reproduction, on survival and on both.

## 3 Selection on reproduction

If selection only acts on reproduction, the mutant produces *r* times more offspring compared to the wild type. The transition probability from state

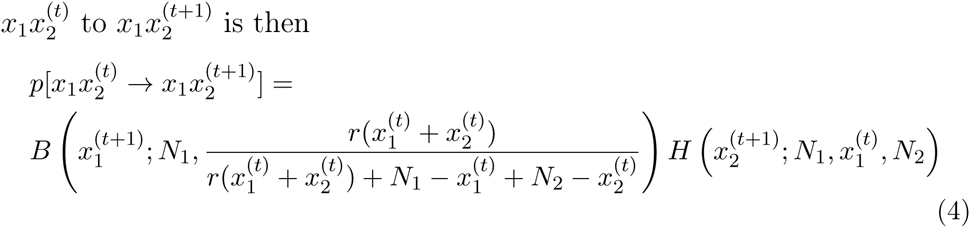

It seems difficult to find a closed form of the fixation probability without making additional assumptions, such as approximating the noncentral hyper-geometric distribution, which arises from biased sampling, with a binomial distribution, which requires that *N*_*i*+1_ is either very small or close to *N*_*i*_ (Emigh, 1979a,b). However, we can obtain some first insights from calculating 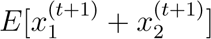, i.e. the expected total number of mutants in the next time step.

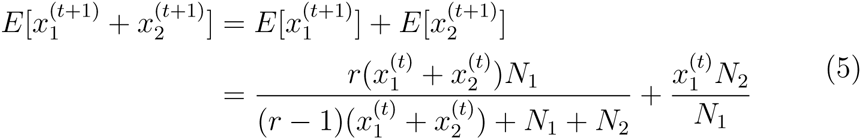

Although 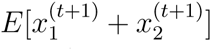 is not a proxy of the fixation probability per se, it illustrates an important aspect of the fixation dynamics.

Given that the total size of the population is fixed, the state of the population is determined by the relative numbers of young and old individuals. Replacing *N*_2_ by *N* − *N*_1_, we observe a minimum of the expected number of mutants in the next time step, where 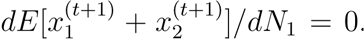. The corresponding number of young individuals is

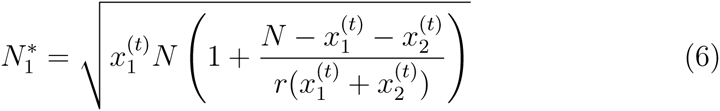

As the expected number of mutants in the next time step depends on the number of young individuals in a non trivial way once we depart from neutrality, the fixation probability of the mutant could in principle also have an extremum in populations with an intermediate fraction of young individuals. A numerical consideration of fixation probabilities shows that the fixation probability indeed has a minimum for an intermediate fraction of young individuals, as shown for a population of total size 20 in Figure 3. In Appendix B we show that also a weak selection analysis recovers the U-shaped pattern of fixation probability, although there are foreseeable disagreements on the exact values, especially when the selective advantage of the mutant becomes large.

**Figure 3:**
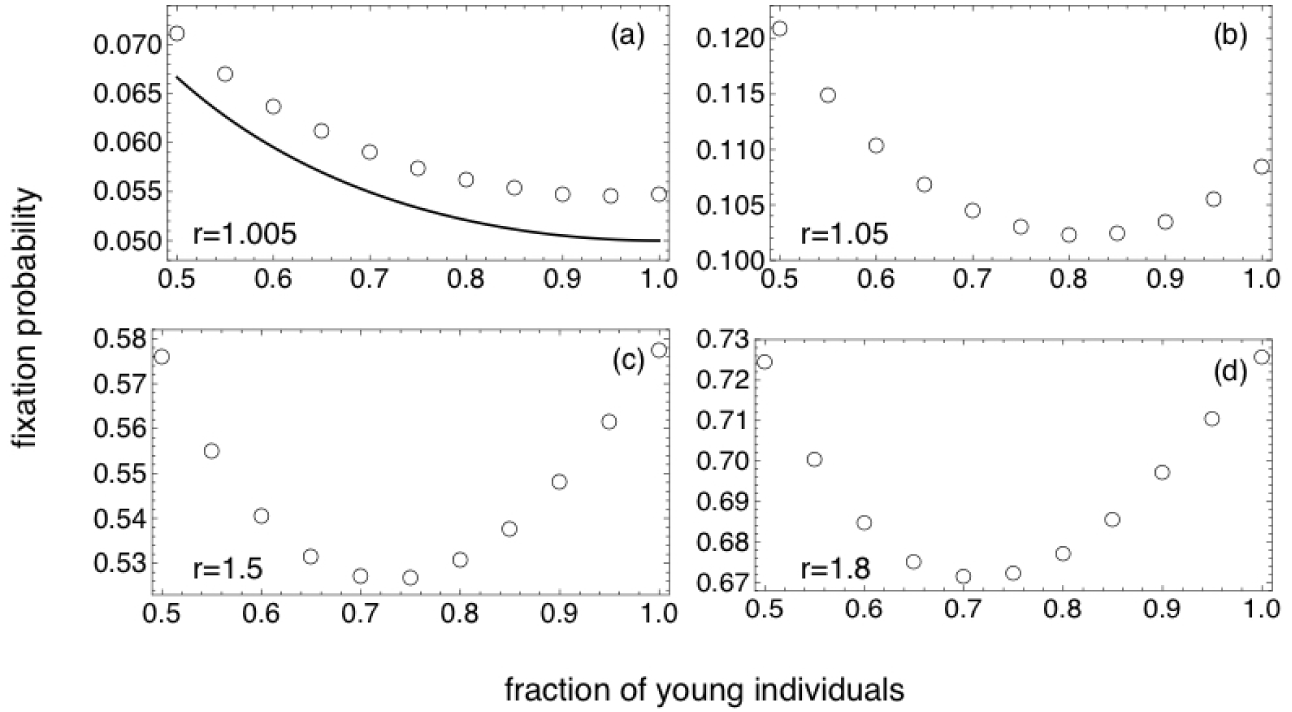
Constant selection on reproduction in populations with fixed total size of 20. Symbols represent the fixation probability of a single young mutant with fitness *r*. The solid line in panel (a) represents the fixation probability in the neutral case when *r* = 1. There is always a minimum of the fixation probability for intermediate fraction of young individuals. The fraction of young individuals at the minimum decreases when *r* increases (panels (b)-(d)). A Mathematica notebook file for generating this figure can be found in the Supplementary Information.

## 4 Selection on survival

If selection only works on survival, the mutant type has the same fecundity as the wild type, but is *r* times more likely to survive to age two. Because of the selective advantage of the mutant, the survival step follows the Wallenius’ noncentral hypergeometric distribution instead of the standard hypergeometric distribution (Fog, 2008b). The stochastic process can be illustrated conveniently with a urn model. Think of a urn with black and white balls. We take *n* balls one by one from the urn without replacement. Every time, a black ball is *r* times more likely to be chosen compared to a white ball. The Wallenius’ noncentral hypergeometric distribution tells us the probability of obtaining *x* black balls by the end of the experiment. Similarly, in our case, there are *N*_1_ mutants and wild-type individuals in the first age class. Among them, *N*_2_ will be chosen to form the next age class. At each draw, a mutant is *r* times more likely to be chosen than a wild type. The Wallenius’ noncentral hypergeometri distribution tells us the probability of obtaining *x*_2_ mutants in the next time step.

Therefore, the transition probability from state 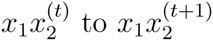 is

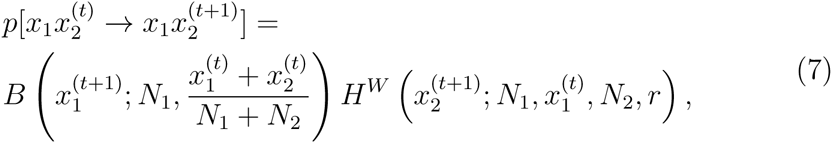
in which *H*^*W*^ is Wallenius’ noncentral hypergeometric distribution, for which closed form implementations are numerically cumbersome (Fog, 2008a).

When the selective advantage of the mutant *r* is small, *H*^*W*^ can be approximated by the corresponding standard hypergeometric distribution, and thus the fixation probability of the mutant is similar to that of the neutral case (Figure 4a). However, when *r* is large, we numerically observe a remarkable increase of the fixation probability in populations with an intermediate fraction of young individuals (Figure 4b-d).

**Figure 4:**
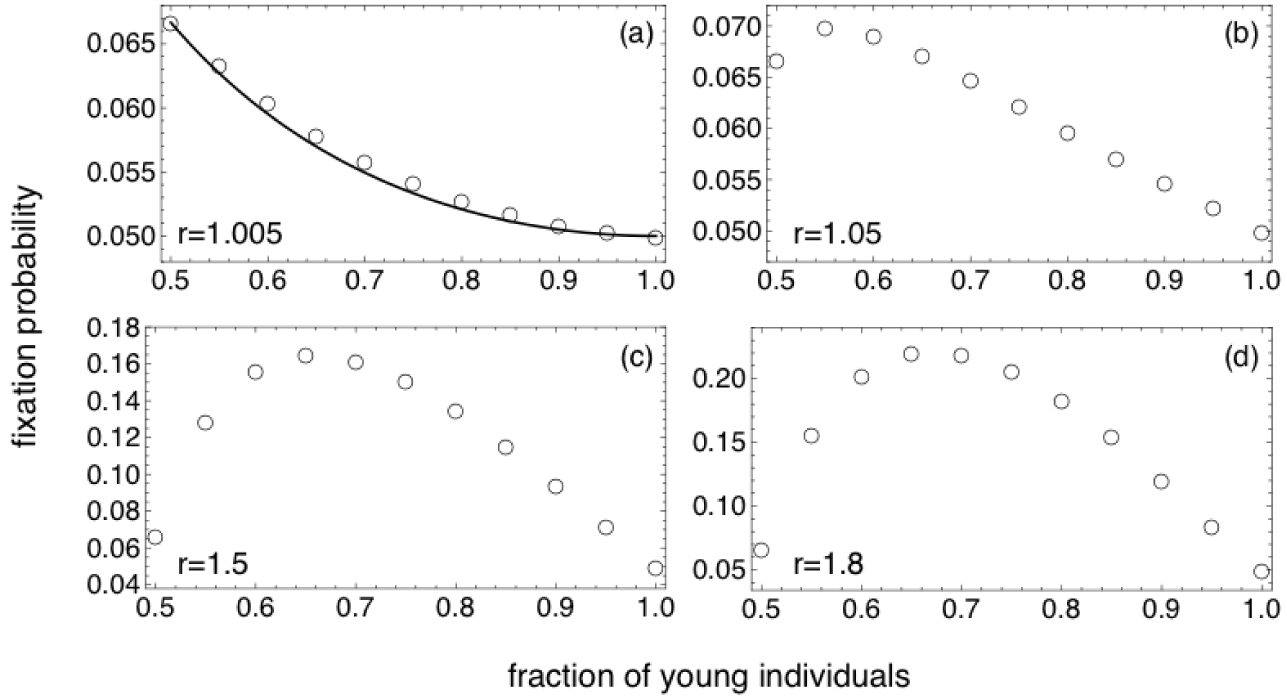
Constant selection on survival in populations with a total fixed size of 20. Symbols represent the fixation probability of one single mutant in the young age class with fitness *r*. The solid line in panel (a) represents the fixation probability in the neutral case, when *r* = 1. When *r* is small, the fixation probability decreases monotonically, resembling the neutral case. But when *r* becomes larger (panels (b)-(d)), there is an intermediate maximum of fixation probability. The corresponding fraction of young individuals increase when *r* increases. A Mathematica notebook file for generating this figure can be found in the Supplementary Information.

Although we cannot calculate the exact fixation probability analytically due to the complexity associated with the Wallenius’ noncentral hypergeometric distribution, it is still possible to understand the pattern of having a maximum of fixation probability in populations with intermediate fraction of young individuals intuitively. First, we consider the change in the expected number of mutants in the young age class, as the fraction of young individuals increase. We keep in mind that selection works on survival and there is no selection on reproduction. Therefore, the expected number of young mutants in the next time step is simply proportional to the global fraction of mutants in the whole population. As *N*_1_ increases by 1, the expected number of mutants in the young age class in the next time step increases by 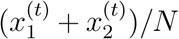, which is between 0 and 1. This is irrespective of the value of *N*_1_.

Next, we consider the change in the expected number of mutants in the old age class, as the fraction of young individuals increase. In the limit of very large selective advantage *r*, basically all mutants in the young age class will be selected to survive into the old age class, if there is enough space, namely, *N*_2_ is greater than the total number of mutants. Otherwise, among all the mutants in the young age class, *N*_2_ of them will survive and enter the old age class. Therefore, the expected number of mutants in the old age class in the next time step is the minimum between the current number of mutants in the young age class, and the total number of old individuals in the population. In short, if *r* ≫ 1, 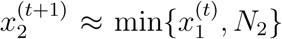. When the fraction of young individual is small, 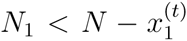, the expected number of mutants in the old age class in the next time step does not change as *N*_1_ increase by one, as each of these mutants will survive. But when the number of young individual is large, 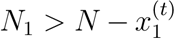, the expected number of mutants in the old age class is limited by *N*_2_. In this case, as *N*_1_ increases by 1, *N*_2_ and thus 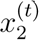 must decrease by one due to the constant population size.

Taken together, the change in the expected total number of mutants in one time step is the sum of the change in the expected number of mutants in the young age class and the expected number of mutants in the old age class. As *N*_1_ increases by one, on the one hand, the expected number of mutants in the young age class in the next step should increase between 0 and 1. On the other hand, the expected number of mutants in the old age class does not change when *N*_1_ is small, but decrease by 1 when *N*_1_ becomes large. Therefore, the expected total number of mutants in the next time step should first increase and then decrease, as *N*_1_ increases. The corresponding fixation probabilities are explored numerically in Fig. 4.

## 5 Selection on both reproduction and survival

In the two previous sections, we have shown that if selection only acts on survival, there is a minimum of the fixation probability when the number of young individuals is intermediate. But if selection acts on survival and when the selective advantage of the mutant is sufficiently large, there is a maximum of the fixation probability when the number of young individuals is intermediate.

If selection works on both reproduction and survival, we assume that a beneficial mutant may not only produce more offspring, but also is more likely to survive to the next age. This double effects could result from the fact that the mutant allocates extra payoff to both reproduction and survival. Consider a mutant that has increased access to food resources compared to the wild type. As a result, she may choose to (or be genetically programmed to) consume the extra food immediately, thereby allocating most of the benefits to reproduction and relatively little to improving her chance of surviving to the next round of reproduction. On the other hand, she may choose to save most of the food for provision, thereby has little improvement in the current round of reproduction, but substantially improvement in the chance of survival. There exists a well established body of allocation theory, with different functional forms for allocating limited resource to reproduction and survival. This further leads to different patterns of aging in different species and populations. For a recent review, see Baudisch and Vaupel (2012). In the case where selection acts on both reproduction and survival, and the mutant allocates its payoff benefit to both, the transition probability from state 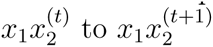 is

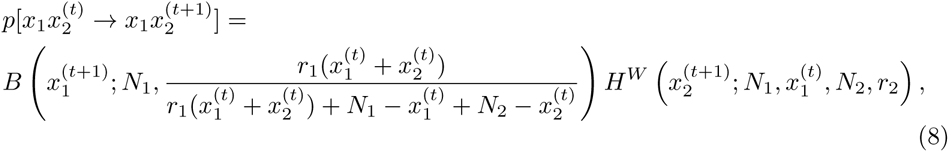
in which *r*_1_ and *r*_2_ represent that the mutant produces *r*_1_ times offspring compared to the wild type, and is *r*_2_ times more likely to survive to the next age.

In this study we focus on demonstrating the distinct effects of different selective forces on the fixation probability of mutants. Therefore, we choose arbitrarily the simple case where *r*_1_ = *r*_2_ to study the model, as Altrock and Traulsen (2009) did in studying the evolutionary dynamics of stochastic birth-death processes. In Appendix C, we show examples of allocating fitness benefits between reproduction and survival following a linear pattern, where *r*_1_ and *r*_2_ are different. Following the same method, the effects of allocating benefits in other ways can be studied. We also show in Appendix C that mutants taking a different life history trade-off strategy also have complex nonlinear patterns of fixation probabilities, depending on the population demographic structure. We provide an example, showing that trading for higher reproduction at the cost of reduced survival produces very different patterns of fixation probability from the other way round. Considering the complexity of this question and the vast diversity in the range of life history trade-off strategies, it is better to study this in detail in a separated work.

From the numerical calculation results (for the special case that *r*_1_ = *r*_2_ = *r*) show in Figure 5, we see the pattern of fixation probability of a single mutant in the young age class has combined effects from both selecting on reproduction and on survival. On one hand, the value of the fixation probability is more similar to the case that selection works solely on reproduction (Note that the end points in the first and the third panel are identical, as in these points the effect of age structure disappears.) On the other hand, the pattern of having an apparent maximum of the fixation probability in populations with intermediate fraction of young individuals, is more similar to the case when selection works solely on survival.

**Figure 5:**
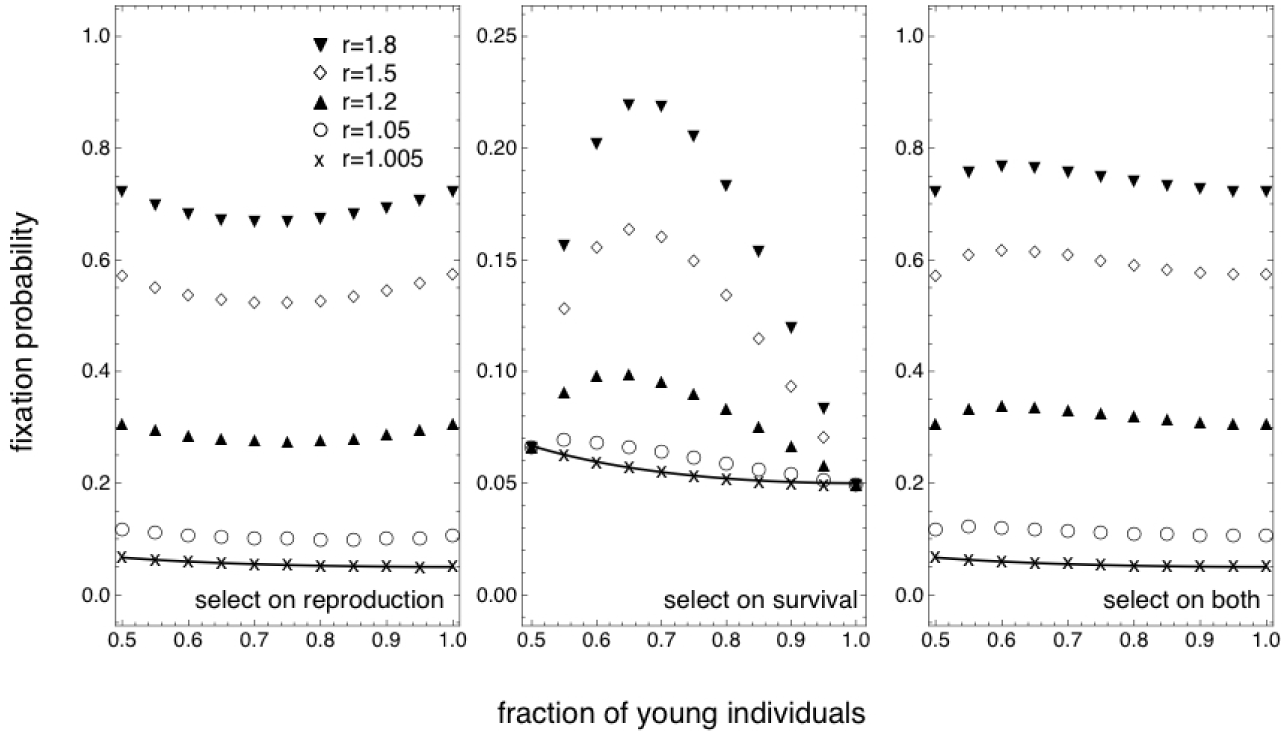
A comparison between selecting on reproduction, survival and both. The figure shows the fixation probability of a single beneficial mutant in the young age class. Solid lines represent the fixation probability in the neutral case, when *r* = 1. The pattern of selecting on both reproduction and survival shows a combined effect. The magnitude of the fixation probability across different age structures is closer to the case when selecting solely on reproduction, but the shape of having an intermediate maximum when *r* is large is similar to selecting on survival (note the different scale on the *y*-axis). Total population size *N* = 20. Mathematica notebook files for generating this figure can be found in the Supplementary Information.

## 6 Effects of population size

Up to now we have shown the effects of demographic population structure on the fixation probability of mutants. It is important to study how the magnitude of the effects scale with the strength of selection and population size. In the following, we show that the relative importance of population size and demographic population structure depends crucially on the strength of selection, in this case, it depends on the relative fitness of the mutant compared to the wild type.

If the relative fitness of the mutant is small (e.g. *r* = 1.005, shown in Figure 6, a-c), population size has a larger effect, and it does not matter too much if selection works on reproduction, survival or both. Because in this limit, instead of demographic structure, the size of population plays the most important role in determining the fixation probability. Fixation or extinction are mainly driven by drift. Therefore the general pattern is that the fixation probability decreases with population size. But if the fitness of the mutant is large (e.g. *r* = 1.2, shown in Figure 6, d-f), the demographic structure of the population plays an essential role. Note that symbols of different colours represent different population sizes. In this case, the symbols correspond to population size 80 and 100 almost overlap, for selecting on reproduction, survival and both. Therefore, the effect of demographic structure on the fixation probabilities does not diminish with population size, but there is convergence for large populations to a certain effect size. The pattern of having an intermediate minimum of fixation probability when selecting on reproduction is preserved irrespective to population size. The intermediate maximum of the fixation probability when selecting on survival exists when the relative fitness of the mutant is not too small.

**Figure 6:**
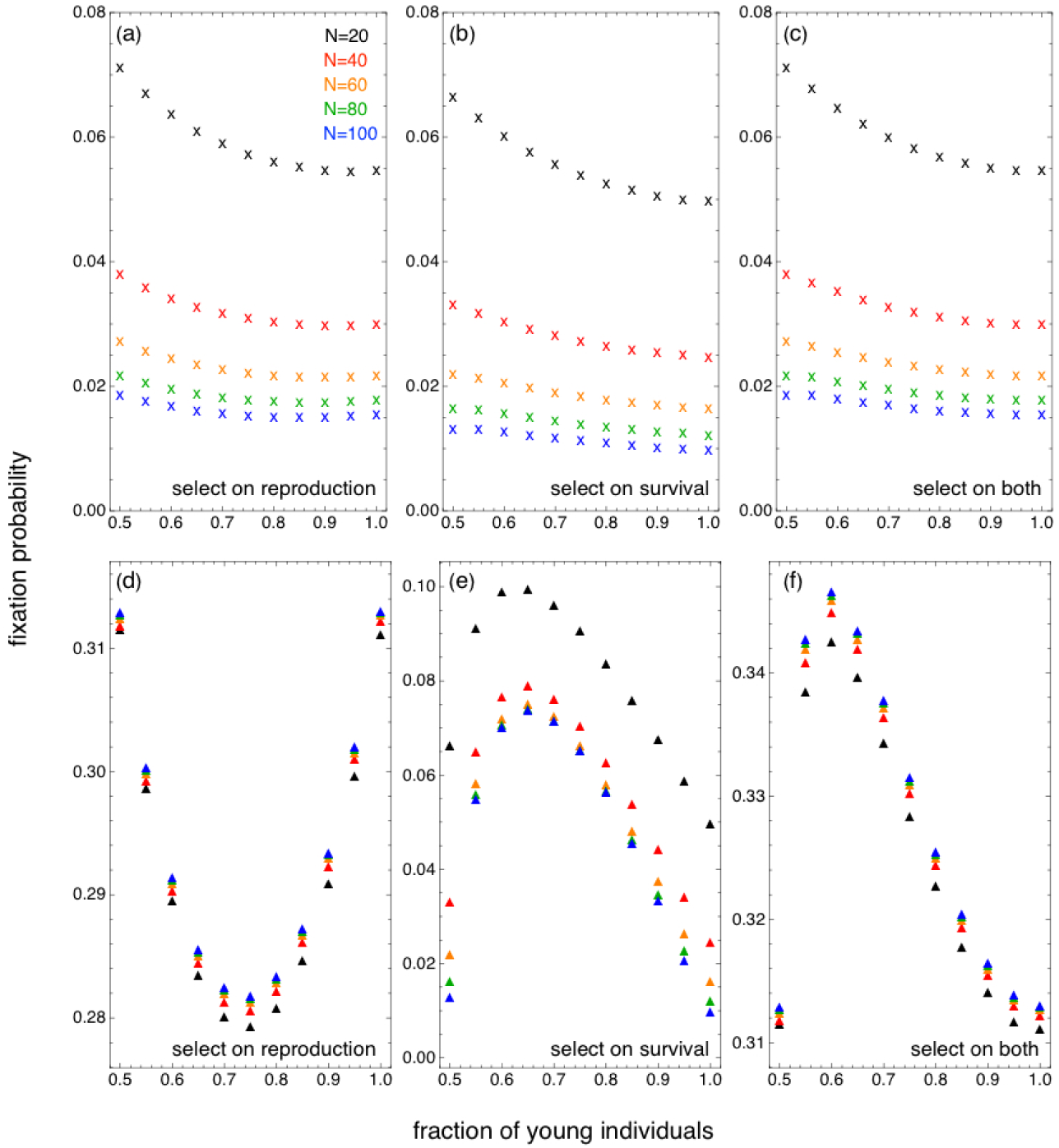
(a-c) Population size has a larger effect on the fixation probability if the relative fitness advantage of the mutant is small, in this case, *r* = 1.005. (d-f) Demographic population structure has a larger effect on the fixation probability if the relative fitness advantage of the mutant is large, in this case, *r* = 1.2. Symbols of different colours represent different population size. Color coding is consistent in all panels. Mathematica notebook files for generating this figure can be found in the Supplementary Information.

Another interesting observation is that the fixation probability of a single mutant is higher in larger populations compared to smaller populations, when *r* is large, and when selecting on reproduction or both reproduction and survival. This is similar to a feature of the Wright-Fisher process (Details in Appendix D).

We also assume that population size is fixed over time. It would be interesting also to study how population size fluctuation may change the dynamics. In general, although the Markov approach we used in the manuscript allows us to calculate the exact fixation probability and it has lead us to find the interesting non-monotonic patterns, it does have limitations in numerically handling large or analysing fluctuating population sizes. In that case we need to update the population state transition matrix dynamically, depending on the detailed population updating processes. Every time the population change its size, the state space is updated. Consequently, the transition probabilities from one state to another also have to change. Although the fixation probability can still be obtained from simulations, it would be challenging to derive analytical results. Compared to the Markov approach, the branching process approach and the diffusion approximation approach may be more suitable modelling frameworks for analysing the effects of population size fluctuation. See Patwa and Wahl (2008) and Wahl (2011) for comprehensive reviews on different approaches of studying the fixation probability of mutants.

In Appendix B, the results we derived from weak selection analysis only require the assumption that the resident population has size *N* when the initial mutant is first introduced, but they do not require this census size to be kept constant thereafter. The diffusion approximation is meant to work when the population size fluctuates over time, despite having unity geometric growth in the limit of infinite size. In populations of finite size, as a result of demographic stochasticity, the total numbers of individuals fluctuate with time. The smaller the population size, the greater the fluctuations. This is because random independent variation in survival and reproduction between individuals is hard to average out when population size is very small (Vindenes et al., 2009). This suggests, qualitatively, the result of having a U-shaped pattern of fixation probability when selecting on reproduction should not be restricted to the situations in which the population size is kept constant over time.

Besides the classic works of Ewens (1967); Kimura and Ohta (1974); Crow (1979), and Otto and Whitlock (1997), there are a number of recent works that study the effects of population size fluctuation on the fixation probability of mutants, including Parsons and Quince (2007a), Orr and Unckless (2008), Engen et al. (2009a,b), Parsons et al. (2010), Uecker and Hermisson (2011) and Waxman (2011). Important insights from the recent developments include the great importance to study the detailed processes of population dynamics, and to distinguish and examine the effects of selection, drift, and the interactions of both. For example, Waxman (2011) pointed out that the changes in population size are not equivalent to the corresponding changes in selection, for it can result in less drift than anticipated. Uecker and Hermisson (2011) showed that even for the same logistic growth of the population, depending on whether it results from the reduction of fertility while keeping mortality constant or the increase of mortality while keeping fertility constant, the fixation probability can be very different, due to stronger effects of drift in the second scenario.

## 7 Beyond two age classes

So far, we have only considered the case of two age classes. In this section, we give a proof-of-principle that the results shown in previous sections are not artifacts of the assumption of two age classes. Instead, the non-trivial effects of population demographic structure on the fixation probability of a mutant are preserved also in populations with multiple age classes.

In a demographically stationary population with *ω* age classes and a total size *N*, assuming the average survival probability *γ* ∈ [0,1] is constant from one age class to the next, the number of individuals in each age class is determined by the equations

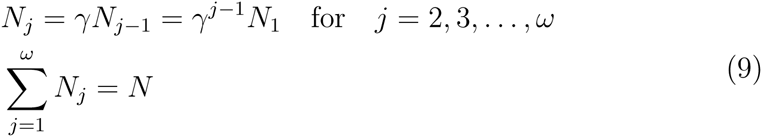

As we are only considering integer *N*_*j*_, the number of individuals in each age class has to be rounded. We do this by calculating the *N*_*j*_ under the condition of a fixed integer *N* first and perform the rounding afterwards. We only show simulations where the *N*_*j*_ add up to our choice of *N*.

The choice of a constant *γ* corresponds to species such as the freshwater hydra (*Hydra magnipapillata* and *Hydra vulgaris*) living under controlled lab conditions (Martínez, 1998; Schaible et al., 2015). In field studies, approximately constant survival probability after the age of reproductive maturity has also been observed in birds, frogs, invertebrates and plants (Baudisch et al., 2013; Jones et al., 2014).

For other species, the survival (mortality) rate can differ widely over different ages or life stages. For example, most mammals including humans have decreasing survival rates after adulthood (*γ* decreases with age). On the other hand, for the desert tortoise the survival probability increases monotonically with age; see Jones et al. (2014) for a review of the diverse patterns of age/life-stage dependent mortality rates.

We show in Figure 7 the patterns of fixation probability of a single mutant from the first age class in a population of three age classes with constant *γ* (*m* = 3). When selection is absent, the fixation probability of the mutant decreases monotonically as the survival rate *γ* decreases. In the boundary case where *γ* = 0, all individuals in the population are in the first age class. Therefore *N*_1_ = *N*, and the fixation probability of the mutant reduces to 1/*N*. In the case where selection works on reproduction or survival, we show that the intermediate minimum or maximum of the fixation probability still persists.

**Figure 7:**
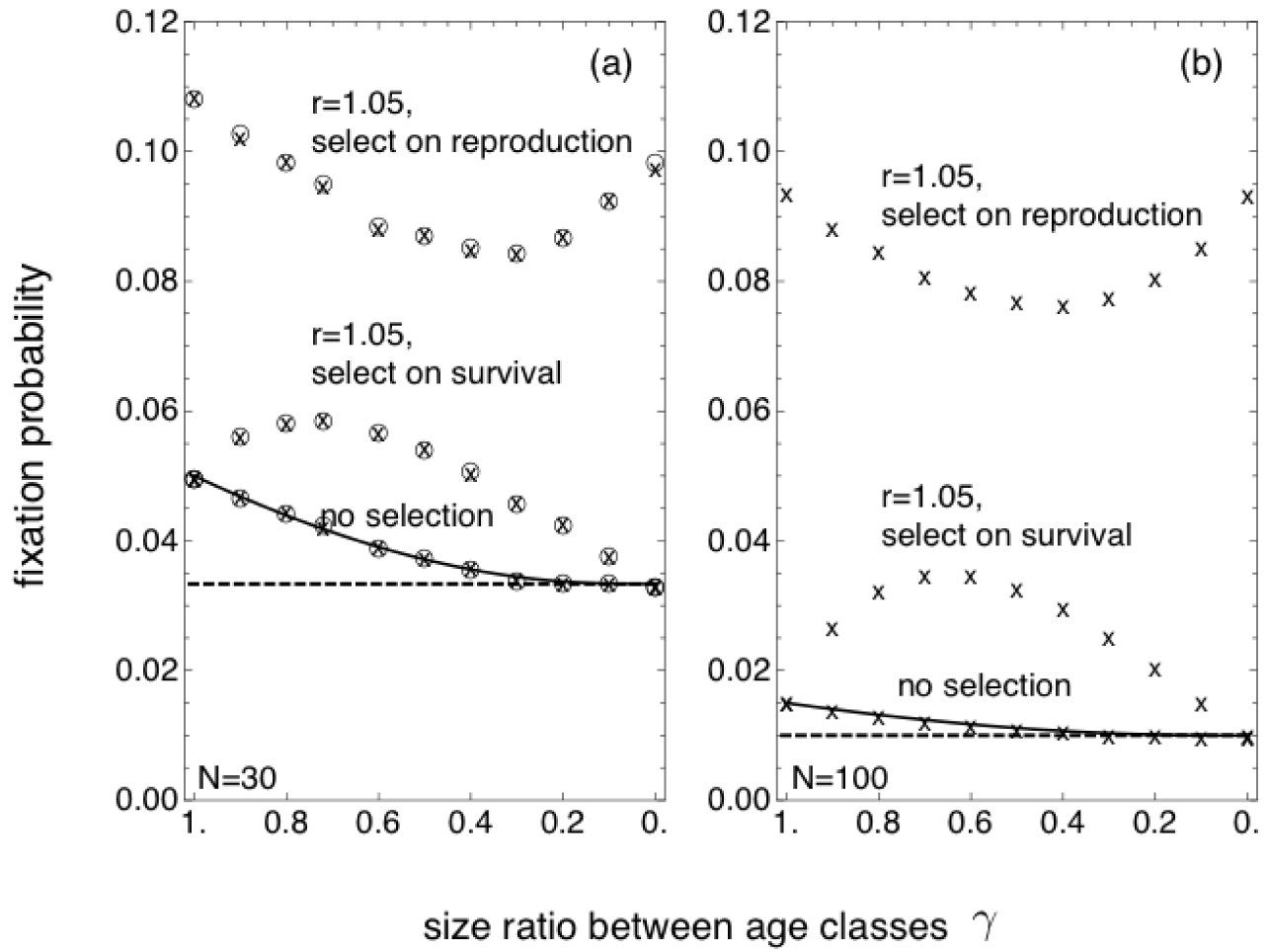
Fixation probability of a single mutant from the first age class in a population of three age classes, depending on the size ratio between age classes, 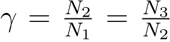. Circles and crosses represent results from numerical solution (by calculating the corresponding eigenvector of the eigenvalue 1 of the population state transition matrix) and simulation, respectively. The solid line represents theanalytical result (Details in Appendix A). Each of the data point in the simulation results is calculated from 10^7^ realizations. The dotted line marks 1/*N*, the fixation probability of a single mutant in a well-mixed age-homogeneous population when selection is absent. (a) For a population size of *N* = 30, we show results from both numerical solution and simulations. The slight wriggles of the fixation probability curves are due to rounding errors when constructing three age classes. These diminish when population size increases. (b) For *N* = 100, we only show results from simulations, because it is computationally costly to calculate the eigenvectors for large matrices.

## 8 Discussion and conclusion

Although it is clear that the likelihood of incorporating a beneficial mutation in the gene pool is a function of the life history architecture of the population, little has been done to pinpoint the direct effects of different life history patterns on promoting or hindering adaptive evolution. In this work, we approach this problem building on the basis of the seminal Felsenstein model of populations with overlapping generations (Felsenstein, 1971). We explore the effects of different sources of evolutionary forces, including selecting on reproduction, survival and a combination of both numerically. We also study the relative importance of the size and demographic structure of the population.

First, our work reveals fundamental differences between selection on reproduction and survival in populations with demographic structures. The differences are remarkable even in the simple case where the population is spatially well-mixed, and neither density nor frequency affects the relative fitness of the mutant. It is known that in well-mixed populations under constant selection, selecting on reproduction or survival is equivalent when mutation is absent (Ewens, 2004; Kaiping et al., 2014). In spatial structured populations, selecting on reproduction and survival are different in general, but it is possible to produce the same effects (Zukewich et al., 2013; Kaveh et al., 2015; Hindersin and Traulsen, 2015). For example, under the Moran process scheme, selecting on reproduction with a birth-death updating rule is still equivalent to selecting on survival with a death-birth updating rule, in any populations with “homogeneous” structures (e.g., lattices, cycles, and island models) and symmetric dispersal (Taylor et al., 2011). It is worthwhile to note that in degree-heterogeneous graphs, the fixation probability of an advantageous mutant depends crucially on if selection works on reproduction or survival (Antal et al., 2006; Hindersin and Traulsen, 2015; Kaveh et al., 2015). Even when selection works only on reproduction, the sequence of birth and death events can lead to completely different evolutionary dynamics (Zukewich et al., 2013; Kaveh et al., 2015). For almost any random graph, a selectively advantageous mutant almost always has higher-than-neutral fixation probability if birth takes places before death, but has lower-than-neutral fixation probability if death happens first (Hindersin and Traulsen, 2015). Although the differences of selecting on reproduction and survival is relatively well recognized and studied in spatially structured populations, less is known in populations with demographic structure.

Even under constant selective advantage, life history of individuals in the population makes the evolutionary dynamics differ drastically with different sources of selective forces (Caswell, 2001; Houston and McNamara, 1999). In the future, it would be interesting to go beyond the simple case of constant selection and investigate the effects of density and frequency dependent fitness of the mutants. In the well-mixed case, low density favors individuals who direct their efforts towards exploring the ecological environment and thereby maximising their reproduction rates. But under high density conditions, since most resources have already been absorbed into the population, it is more efficient to direct one’s effort towards exploiting other population members through interactions (Blute, 2011). Usually, the successfulness of a strategy is not determined by its nature alone, but on the presence and frequency of other strategies in the population. Density and frequency effects are hardly disconnected from each other in natural populations, and it is often necessary to take both into account in order to capture the important features of evolutionary dynamics, especially when there is substantial change in population size (Novak et al., 2013; Li et al., 2015b; Huang et al., 2015). Under the framework of Evolutionary Game Theory, elegant conditions such as the 1/3 rule have been obtained (Nowak et al., 2004; Imhof and Nowak, 2006; Lessard and Ladret, 2007), which determine if strategies such as cooperation can evolve in the first place. But it is unclear if such conditions still hold in populations with demographic structures. Therefore, it seems interesting to study the coevolution between population demography and game theoretic strategies in future work.

In natural populations, species display an enormous variety of different life history patterns (Jones et al., 2014). Often mutations affect the fitness of individuals in both reproductive and survival aspects. A classical example is the throughly discussed ornamented trains of male peacocks, which are sublimely beneficial in terms of mating success, but at the same time tremendously costly when it comes to the chance of escaping from predators. Another beautiful example is the shape of wings in migrating birds. Pointed wings are aerodynamically desirable for fast and long-distance flights (Mönkkönen, 1995; Berthold, 1996; Hedenström, 2002; Bowlin and Wikelski, 2008; Minias et al., 2015). But on the other hand, they reduce the manoeuvrability that help birds with foraging and courtship displays (Alatalo et al., 1984; Swaddle and Lockwood, 2003). Precisely due to the high benefits and high costs of pointed wings, the change of selective pressure leads to the change of wing morphology in bird populations. This has been observed in many passerine species, e.g. the fast evolution of stonechats Saxicola torquata in response to changing environmental conditions (Baldwin et al., 2010). It is interesting also to note that evolution can go to great detail in balancing the cost and benefits of a single morphological trait, and life history serves as a channel through which evolutionary forces fine tune the balance. As an example and also a demonstration of its plastic potential, it is found that the juveniles of migrating birds have less pointed wings compared to adults. This could be due to the fact that juveniles are more naive and thus more vulnerable to predators. Under elevated predation pressure, improved manoeuvrability is more important than migration performance, particularly in the early stage of life (Pérez-Tris and Tellería, 2001). For more examples of the amazing variety in survival and reproduction trajectories over the life course across species, Jones et al. (2014) provides a recent review.

In this work, we focus mostly on the simple case of two age classes. But it is possible to extend our results to cases with multiple age classes, as demonstrated in Section 7. We have shown how the fixation probability of a mutation with effect on certain components of the life history is influenced by the demographic structure of the resident population, i.e. the relative fraction of young versus adult individuals. We note that a natural interpretation of the same results can also be given in terms of the rate of aging that characterizes the resident population. In fact, the relative abundances of young and adult individuals depend on the probability of newborn survival to young age and the probability of surviving from young age to old age. Aging is defined as an age related deterioration in survival. Aging features in a large number of species and is one of the most salient life history traits. It follows from the definition of aging that, in our model, populations that are composed by a higher fraction of young individuals are characterized by a higher rate of aging (i.e. stronger deterioration in survival with age). Conversely, populations with a higher representation of old individuals may possess no aging at all or even negative aging (i.e. survival does not decline but it may even increase with age, see Baudisch and Vaupel (2012) and Jones et al. (2014)).

In this way, our results can be directly linked with the effect that an important life history trait exerts on adaptive evolution. If natural selection acts on reproduction, for any beneficial mutant, intermediate rates of aging always reduce the fixation probability, although the chance of fixation may be higher in populations with low or negative rates of aging, depending on the difference in fitness between the mutant and the wild type. If natural selection acts on survival and the fitness difference between the beneficial mutant and the wild type is not too small, there is a maximum of fixation probability in populations with intermediate rates of aging, while in populations with extreme positive or negative rates of aging, the fixation probability of the beneficial mutant is reduced. In this regard, it would be of interest to study variation in the rate of adaptation in those systems that have already been established for within-species comparison of the rate of aging across populations. For example, populations of guppies living under different environmental conditions have proven to be a valuable model to understand ecologically-induced variation in aging within a single species (Reznick et al., 2004; Bronikowski and Promislow, 2005). Similarly, different life-history ecotypes of the garter snake show different aging patterns (Sparkman et al., 2007; Robert and Bronikowski, 2010). If natural selection acts on both survival and reproduction, depending on how fitness advantages are allocated and the magnitude of the fitness difference between mutant and the wild type, complex patterns with multiple extrema of the fixation probability with respect to the rate of aging can emerge.

## 9 Summary

To summarize, in this work we have studied the direct effects of population demographic structures on stochastic evolutionary dynamics, under the constant selection regime. Using a model with two age classes and constant population size, we have compared the fixation probability of mutants under different population demographic structures. Different targets of selective forces, as well as the relative impacts of the size and structure of the populations are also evaluated in our analyses. Through this work, we hope to call attention to the importance of considering life history when studying evolutionary dynamics. Facilitated by modern computational power, now we have the opportunity to delve into many interesting questions that were technically difficult to approach a few decades ago. Our work opens up new directions for future research, including the coevolution of population structure, resource allocation and strategic dynamics, the impacts of demography on the rate of adaptive evolution, and the density/frequency dependent fitness effects in populations with different life history patterns.

## Acknowledgments

We thank Sabin Lessard (University of Montreal), whose suggestions have led to substantial improvement of the manuscript. We also thank Marion Blute (University of Toronto), Barbara Helm (University of Glasgow), Yasuo Ihara (University of Tokyo), and Miriam Liedvogel (MPI for Evolutionary Biology) for constructive discussions. X.L. is grateful to the International Max Planck Research School (IMPRS) for Evolutionary Biology for funding and support. Funding for S.G. was provided by the Max Planck International Research Network on Aging (MaxNetAging).

## A Fixation probability of a single mutant in the young age class

Fisher’s reproductive value *v*_*k*_ of an individual in age class *k* in an age-structured population with pre-breeding census is (Charlesworth, 1994):

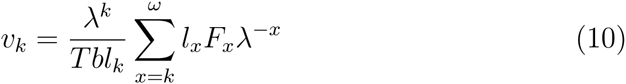
in which *λ* is the asymptotic growth rate of the population, *l*_*k*_ = *S*_1_*S*_2_…*S*_*k*–1_ is the probability of an individual surviving at least to age class *k* with *l*_1_ = 1, *F*_*k*_ is the expected number of offspring to an individual in age class k, *T* is the generation time in the stable population, *b* is the birth rate in the stable population, and *ω* is the maximum attainable age.

In our specific model of two age classes (*ω* = 2), the parameter values are, *λ* = 1 (constant population size), 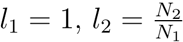, and *F*_1_ = *F*_2_ = *N*_1_/*N*, where *N* = *N*_1_ + *N*_2_.

We have *v*_1_ = 1 and *v*_2_ = *N*_1_/*N*. Following Emigh (1979a), the fixation probability of a selectively neutral mutant is

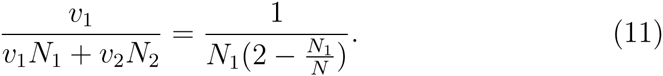

In the case of multiple age classes as in Section 7, *l*_*k*_ = *γ*^*k*–1^, in which *γ* is the survival probability to the next age class, 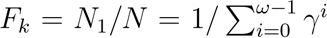 for all *k*. The size of the *k*-th age class is 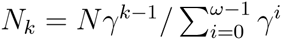. Therefore, the reproductive value of the *k*-th age class in a stationary population is

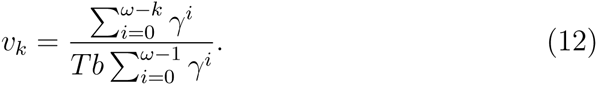

The fixation probability of a single mutant in the first age class is then

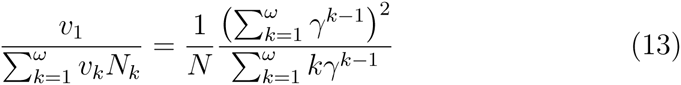

In the case of two age classes, *ω* = 2, *γ* = *N*_2_/*N*_1_. Denote the fraction of young individuals *ƒ* = *N*_1_/*N*, the fixation probability recovers (*Nf*(2 − *ƒ*))^−1^.

## B Weak selection analysis for selecting on reproduction

Following Kimura (1957, 1962); Emigh (1979a,b) and Vindenes et al. (2009), in an age-structured population of haploid individuals, the fixation probability of a single mutant with selective advantage *s* in age class *i* can be approximated by

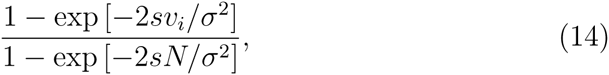
where *N* is the resident population size, *σ*^2^ is the expected variation in geometric growth of the wild-type population, and *v*_*i*_ is the reproductive value of a wild-type individual in the same age class as the initial mutant. The selective advantage *s* of a beneficial mutation is computed as the absolute value of the difference between the stable growth rate of the wild-type, which is the leading eigenvalue of its projection matrix (Caswell, 2001), and the stable growth rate that would be observed in a population composed entirely by mutants (i.e. the leading eigenvalue of the mutant Leslie matrix). Following Engen et al. (2005) in assuming no covariances between matrix elements, the demographic variance *σ*^2^ can be approximated by

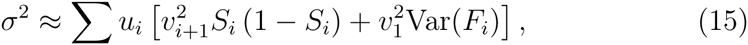

In this expression, *S*_*i*_ is the (*i* + 1)^*th*^ element of column *i* of the (pre-breeding census) Leslie matrix, representing the survival rate of an individual in age class *i*; *F*_*i*_ is the *i*^*th*^ element of the first row of the matrix, representing the numbers of offspring produced by an individual currently in age class *i*; *u*_*i*_ and *v*_*i*_ are the *i*^*th*^ elements of the right and left leading eigenvectors, respectively, of the Leslie matrix and are scaled so that Σ *u*_*i*_ = 1 and Σ *u*_*i*_*v*_*i*_ = 1.

In our model, when selecting on reproduction, a mutant produces *r* times of offspring compared to the wild-type. Hence, the selective advantage *s* of the mutant can be calculated as the absolute value of the difference between the leading eigenvalue of the Leslie matrix of the wild-type (which is unity in our case of constant size) and the leading eigenvalue of the same matrix with the first row multiplied by *r*, that is, the leading eigenvalue of the following matrix,

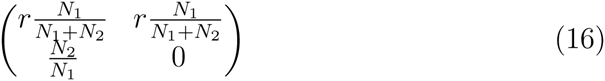

In order to compute the demographic variance, we calculate Var(*F*_*i*_) by treating first row elements of the resident Leslie matrix as averages of Poisson distributions. Figure 8 (corresponding to Figure 3 in the main text) and Figure 9 (corresponding to panels (a) and (d) in Figure 5 in the main text) show the results of using Eq.(14) with the same *r* and *N* values used in our numerical solutions.

**Figure 8:**
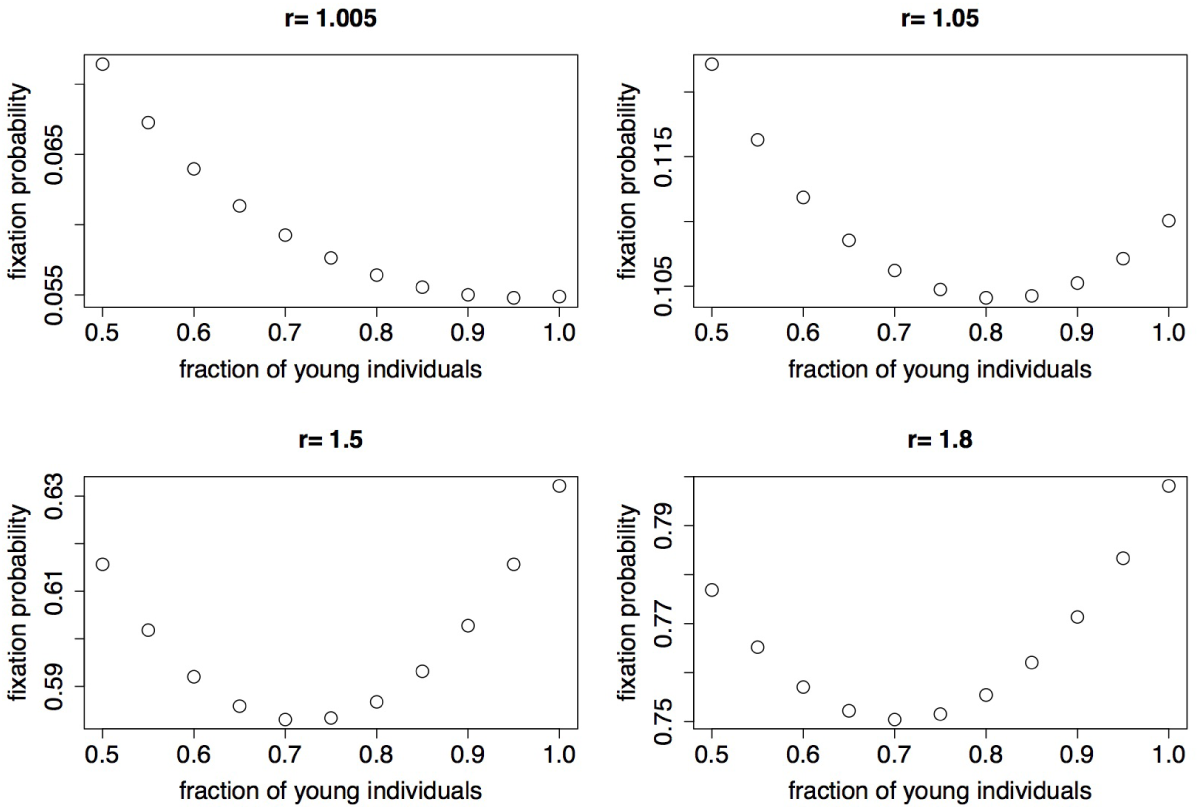
Weak selection approximation to the fixation probability of a mutant with reproductive advantage *r*.

**Figure 9:**
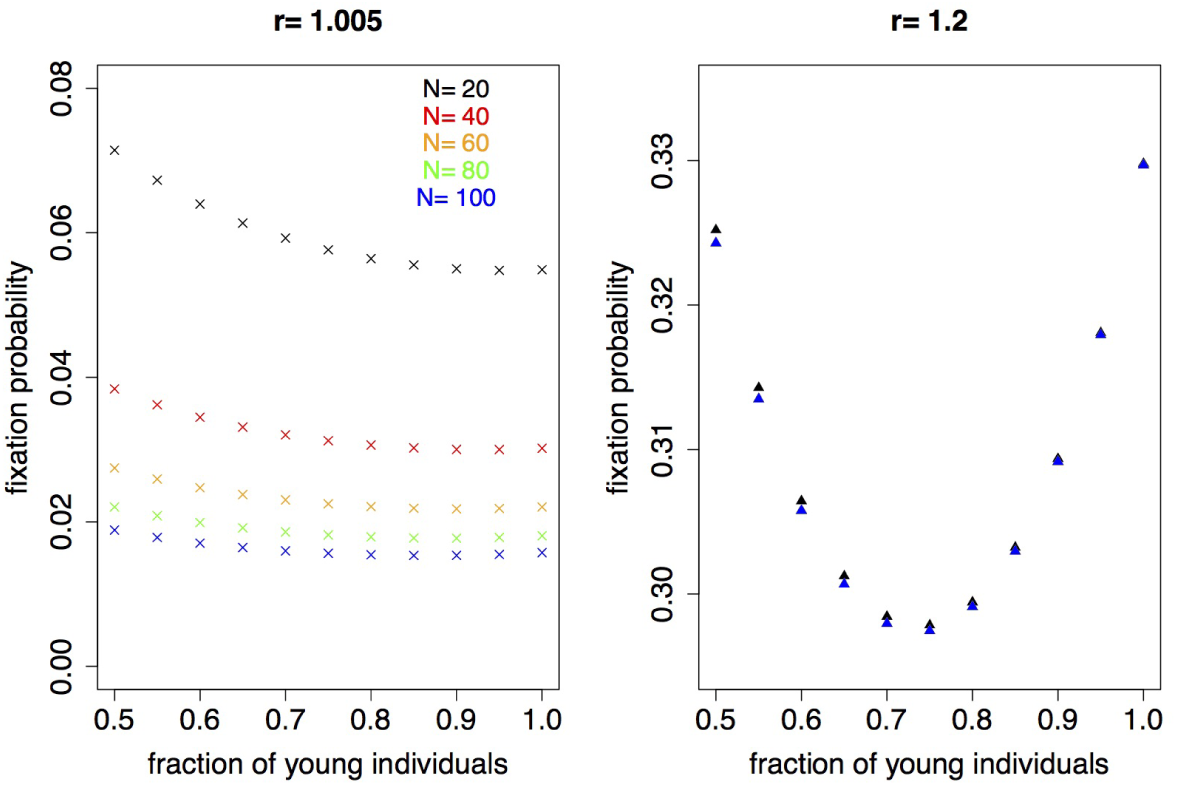
Weak selection approximation to the fixation probability of a mutant with reproductive advantage for different population size and selective advantage. Note that in the right panel, when selective advantage of the mutant is relatively large (*r* = 1.2), the symbols representing different population sizes overlap with each other (especially from N=40 to N=100).

Despite the foreseeable quantitative disagreements especially when the selective advantage becomes large, the weak selection approximation results preserve the U-shaped pattern of fixation probability, under different selection intensities and at different population sizes.

## C Allocation and trade-off of fitness benefits

In Section 5 we show the combined effects of selecting on reproduction and survival at the same time. There we use an arbitrary example of *r*_1_ = *r*_2_ = *r* for simplicity. In this case, the mutant type not only produces *r* times more offspring than the wild type, but also is *r* times more likely to survive to the next age class. This of course is a very special case. It corresponds to the scenario that the mutant happens to split its payoff increment “equally” on improving reproduction and survival. In nature, mutants can in principle allocate the payoff increment in any combination of improving reproduction and survival. The mutant can even commit so much to improving one of them at the cost of reducing the other. This leads to the life history trade-off. Depending on the physiological nature of the species and social interactions in the population, the consequence of putting more weight on one aspect at the cost of reducing the other can be very complex. To illustrate this, in the following we gave examples of allocating an increment of benefit in different ways, and their consequences on the fitness probability of the mutant.

First, imagine that the mutant allocates its extra benefit in a linear way. Relative to wild types, the mutant has an extra payoff *s*. She allocates *a* fraction of it to increase her reproduction, and use the rest to improve survival. Therefore her relative fitness for reproduction *r*_1_ = 1 + *as*, and her relative fitness for survival *r*_2_ = 1 + (1–*a*)*s*. In Figure 10 we show the effects of this particular way of allocating extra benefits on the fixation probability of mutants.

**Figure 10:**
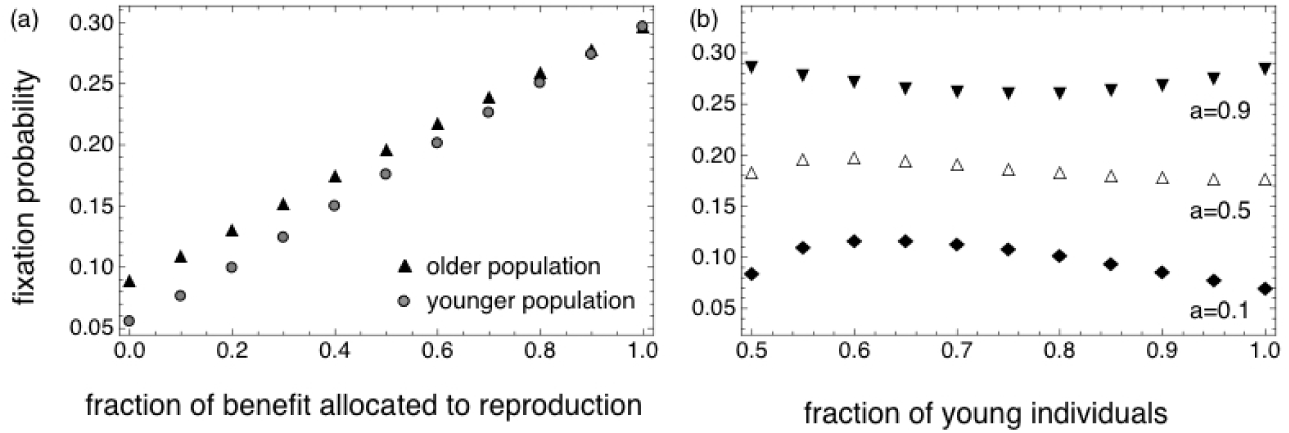
Effects on fixation probability of a mutant that has an extra fitness benefit of *s* = 0.2. (a) The fixation probability increases monotonically with the fraction of benefits allocated to reproduction. In the older population, the fraction of young individual is 0.55, and in the younger population, the fraction of young individual is 0.95. (b) If the fraction of benefits that can be allocated to reproduction is fixed, the fixation probability varies in a non-linear way in populations with different demographic structures. *a* is the fraction of extra benefits that is allocated to reproduction (*N* = 20, a Mathematica notebook file for generating this figure can be found in the Supplementary Information).

It is also very interesting to consider the reproduction/survival trade-off. Considering the great variety in life history and different ways of resource allocation, here we present very briefly an example in Figure 11, that trading on reproduction or trading on survival have very different influences on the fixation probability. Given the apparent complexity in the result, it seems challenging to analyse in its whole breadth.

**Figure 11:**
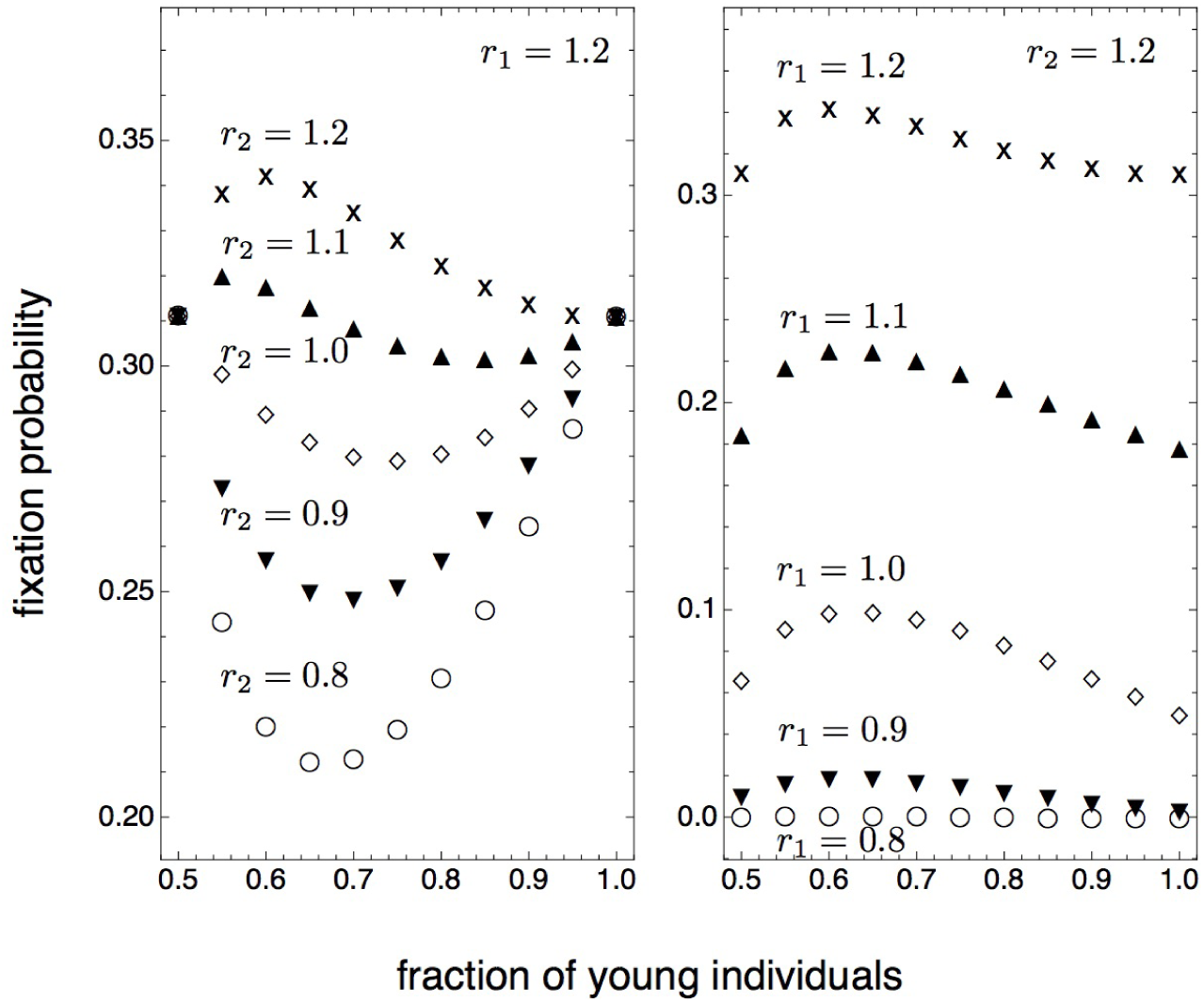
The effects of trading on reproduction or survival. When selection works on both reproduction and survival, a beneficial mutant may have an increase in both of them (the trade-on scenario), or an increase in one of them at the cost of decreasing the other (the trade-off scenario), or the maintenance of the other (zero trade-off scenario). We use *r*_1_ to denote the selective advantage on survival and *r*_2_ to denote the selective advantage on reproduction. (a) Keep selective advantage on reproduction fixed, we change the selective advantage on survival, with two trade-on scenarios, two trade-off scenarios and the zero trade-off scenario. (b) Keep the selective advantage on survival fixed, we change the selective advantage on reproduction, also with two trade-on scenarios, two trade-off scenarios and the zero trade-off scenario.

## D The fixation probability increases with population size in the Wright Fisher process when *r* is large

As we are interested in small populations, we explore the fixation probability in the Wright Fisher process here numerically directly from the associated transition matrix. When *r* is small, the fixation probability of a beneficial mutant decreases with increasing size of the population, approaching 2*s* = 2(*r*–1). But when *r* is large, the fixation probability first decreases and than increase again, although the magnitude of this effect is small, see Figure 12.

**Figure 12:**
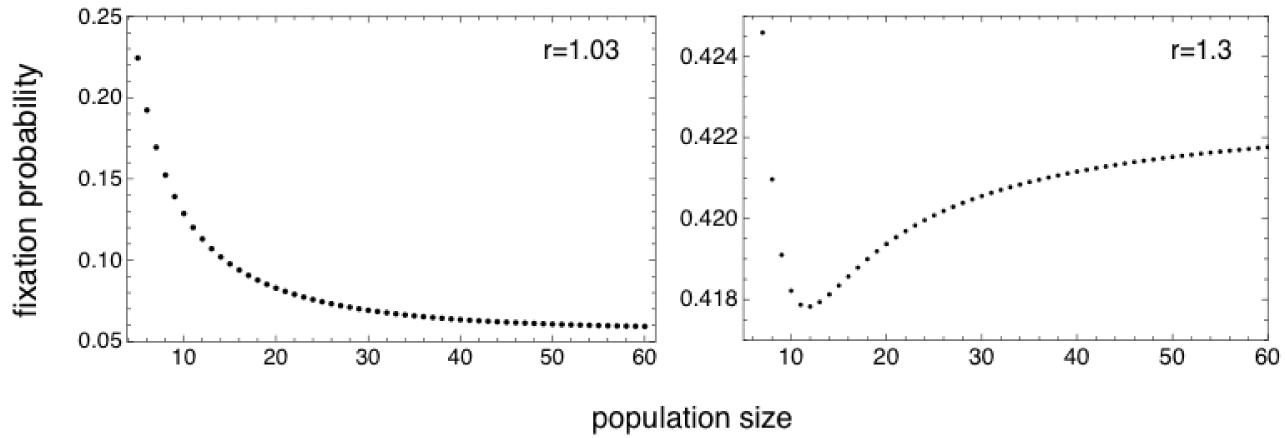
Fixation probability of a single beneficial mutant in the Wright-Fisher process, calculated numerically from the transition matrix. In the left panel when the selective advantage of the mutant is small, as population size grows, the fixation probability approaches the classic result of 2(*r*–1). In the right panel, when the conditions of diffusion approximation are not satisfied, the fixation probability first decreases and then increase again as population size increases (A Mathematica notebook file for generating this figure can be found in the Supplementary Information).

## References

Alatalo, R. V., Gustafsson, L., Lundbkrg, A., 1984. Why do young passerine birds have shorter wings than older birds? Ibis 126, 410–415.

Altrock, P. M., Traulsen, A., 2009. Deterministic evolutionary game dynamics in finite populations. Physical Review E 80, 011909.

Antal, T., Redner, S., Sood, V., 2006. Evolutionary dynamics on degree-heterogeneous graphs. Physical Review Letters 96 (18), 188104.

Baldwin, M. W., Winkler, H., Organ, C. L., Helm, B., 2010. Wing pointedness associated with migratory distance in common garden and comparative studies of stonechats (saxicola torquata). Journal of Evolutionary Biology 23, 1050–1063.

Baudisch, A., Salguero-Gómez, R., Jones, O. R., Wrycza, T., Mbeau-Ache, C., Franco, M., Colchero, F., 2013. The pace and shape of senescence in angiosperms. Journal of Ecology 101, 596–606.

Baudisch, A., Vaupel, J. W., 2012. Getting to the root of aging. Science 338, 618–619.

Berthold, P., 1996. Control of bird migration. Springer Science & Business Media.

Blute, M., 2011. Super cooperators? Trends in Ecology & Evolution 26, 624–625.

Bowlin, M. S., Wikelski, M., 2008. Pointed wings, low wingloading and calm air reduce migratory flight costs in songbirds. PLoS One 3, e2154.

Bronikowski, A. M., Promislow, D. E., 2005. Testing evolutionary theories of aging in wild populations. Trends in Ecology & Evolution 20, 271–273.

Caswell, H., 2001. Matrix population models, 2nd Edition. Sinauer Associates.

Charlesworth, B., 1994. Evolution in age-structured populations, 2nd Edition. Cambridge University Press Cambridge.

Charlesworth, B., 2001. The effect of life-history and mode of inheritance on neutral genetic variability. Genetical Research 77, 153–166.

Crow, J. F., 1979. Gene frequency and fitness change in an age-structured population. Annals of Human Genetics 42, 355–370.

Crow, J. F., Kimura, M., 1970. An Introduction to Population Genetics Theory. Harper and Row, New York.

Emigh, T. H., 1979a. The dynamics of finite haploid populations with overlapping generations. i. moments, fixation probabilities and stationary distributions. Genetics 92, 323–337.

Emigh, T. H., 1979b. The dynamics of finite haploid populations with overlapping generations. ii. the diffusion approximation. Genetics 92, 339–351.

Engen, S., Lande, R., Saether, B.-E., 2005. Effective size of a fluctuating age-structured population. Genetics 170, 941–954.

Engen, S., Lande, R., Sæther, B.-E., 2009a. Fixation probability of beneficial mutations in a fluctuating population. Genetics Research 91, 73.

Engen, S., Lande, R., Sæther, B.-E., 2009b. Reproductive value and fluctuating selection in an age-structured population. Genetics 183, 629–637.

Ewens, W. J., 1967. The probability of survival of a new mutant in a fluctuating enviroment. Heredity 22, 438–443.

Ewens, W. J., 2004. Mathematical Population Genetics. I. Theoretical Introduction. Springer, New York.

Felsenstein, J., 1971. Inbreeding and variance effective numbers in populations with overlapping generations. Genetics 68, 581–597.

Fisher, R. A., 1930. The Genetical Theory of Natural Selection. Clarendon Press, Oxford.

Fog, A., 2008a. Calculation methods for Wallenius’ noncentral hyper-geometric distribution. Communications in Statistics—Simulation and Computation® 37, 258–273.

Fog, A., 2008b. Sampling methods for Wallenius’ and Fisher’s noncentral hypergeometric distributions. Communications in Statistics—Simulation and Computation® 37, 241–257.

Hedenström, A., 2002. Aerodynamics, evolution and ecology of avian flight. Trends in Ecology & Evolution 17, 415–422.

Hindersin, L., Traulsen, A., 2015. Most undirected random graphs are amplifiers of selection for birth-death dynamics, but suppressors of selection for death-birth dynamics. PLoS Computational Biology 11, e1004437.

Houston, A. I., McNamara, J. M., 1999. Models of adaptive behaviour: an approach based on state. Cambridge University Press, Cambridge.

Huang, W., Hauert, C., Traulsen, A., 2015. Stochastic game dynamics under demographic fluctuations. Proceedings of the National Academy of Sciences of the United States of America 112, 9064–9069.

Hubbarde, J. E., Wild, G., Wahl, L. M., 2007. Fixation probabilities when generation times are variable: The burst–death model. Genetics 176, 1703–1712.

Imhof, L. A., Nowak, M. A., 2006. Evolutionary game dynamics in a Wright-Fisher process. Journal of Mathematical Biology 52, 667–681.

Jones, O. R., Scheuerlein, A., Salguero-Gómez, R., Camarda, C. G., Schaible, R., Casper, B. B., Dahlgren, J. P., Ehrlén, J., García, M. B., Menges, E. S., et al., 2014. Diversity of ageing across the tree of life. Nature 505, 169–173.

Kaiping, G., Jacobs, G., Cox, S., Sluckin, T., 2014. Nonequivalence of updating rules in evolutionary games under high mutation rates. Physical Review E 90, 042726.

Kaveh, K., Komarova, N. L., Kohandel, M., 2015. The duality of spatial death-birth and birth-death processes and limitations of the isothermal theorem. Journal of the Royal Society Open Science 2 (140465).

Kimura, M., 1957. Some problems of stochastic processes in genetics. The Annals of Mathematical Statistics 28, 882.

Kimura, M., 1962. On the probability of fixation of mutant genes in a population. Genetics 47, 713–719.

Kimura, M., Ohta, T., 1974. Probability of gene fixation in an expanding finite population. Proceedings of the National Academy of Sciences USA 71, 3377–3379.

Lambert, A., 2006. Probability of fixation under weak selection: a branching process unifying approach. Theoretical population biology 69, 419–441.

Lessard, S., Ladret, V., 2007. The probability of fixation of a single mutant in an exchangeable selection model. Journal of Mathematical Biology 54, 721–744.

Li, X.-Y., Giaimo, S., Baudisch, A., Traulsen, A., 2015a. Modeling evolutionary games in populations with demographic structure. Journal of Theoretical Biology 280, 506–515.

Li, X.-Y., Pietschke, C., Fraune, S., Altrock, P. M., Bosch, T. C., Traulsen, A., 2015b. Which games are growing bacterial populations playing? Journal of The Royal Society Interface 12, 20150121.

Martínez, D. E., 1998. Mortality patterns suggest lack of senescence in hydra. Experimental gerontology 33, 217–225.

Minias, P., Meissner, W., Włodarczyk, R., Ożarowska, A., Piasecka, A., Kaczmarek, K., Janiszewski, T., 2015. Wing shape and migration in shore-birds: a comparative study. Ibis 157, 528–535.

Mönkkönen, M., 1995. Do migrant birds have more pointed wings?: a comparative study. Evolutionary Ecology 9, 520–528.

Novak, S., Chatterjee, K., Nowak, M. A., 2013. Density games. Journal of Theoretical Biology 334, 26–34.

Nowak, M. A., Sasaki, A., Taylor, C., Fudenberg, D., 2004. Emergence of cooperation and evolutionary stability in finite populations. Nature 428, 646–650.

Nowak, M. A., Sigmund, K., 2004. Evolutionary dynamics of biological games. Science 303, 793–799.

Nunney, L., 1991. The influence of age structure and fecundity on effective population size. Proceedings of the Royal Society B: Biological Sciences 246, 71–76.

Nunney, L., 1996. The influence of variation in female fecundity on effective population size. Biological Journal of Linnean Society 59, 411–425.

Orr, H. A., Unckless, R. L., 2008. Population extinction and the genetics of adaptation. The American Naturalist 172, 160–169.

Otto, S. P., Whitlock, M. C., 1997. The probability of fixation in populations of changing size. Genetics 146, 723–733.

Parsons, T. L., Quince, C., 2007a. Fixation in haploid populations exhibiting density dependence i: the non-neutral case. Theoretical population biology 72, 121–135.

Parsons, T. L., Quince, C., 2007b. Fixation in haploid populations exhibiting density dependence ii: The quasi-neutral case. Theoretical population biology 72, 468–479.

Parsons, T. L., Quince, C., Plotkin, J. B., 2010. Some consequences of demographic stochasticity in population genetics. Genetics 185, 1345–1354.

Patwa, Z., Wahl, L. M., 2008. The fixation probability of beneficial mutations. Journal of The Royal Society Interface 5, 1279–1289.

Pérez-Tris, J., Tellería, J. L., 2001. Age-related variation in wing shape of migratory and sedentary blackcaps sylvia atricapilla. Journal of Avian Biology 32, 207–213.

Reznick, D. N., Bryant, M. J., Roff, D., Ghalambor, C. K., Ghalambor, D. E., 2004. Effect of extrinsic mortality on the evolution of senescence in guppies. Nature 431, 1095–1099.

Robert, K. A., Bronikowski, A. M., 2010. Evolution of senescence in nature: physiological evolution in populations of garter snake with divergent life histories. The American Naturalist 175, 147–159.

Schaible, R., Scheuerlein, A., Dańko, M. J., Gampe, J., Martínez, D. E., Vaupel, J. W., 2015. Constant mortality and fertility over age in hydra. Proceedings of the National Academy of Sciences 112, 15701–15706.

Sparkman, A. M., Arnold, S. J., Bronikowski, A. M., 2007. An empirical test of evolutionary theories for reproductive senescence and reproductive effort in the garter snake thamnophis elegans. Proceedings of the Royal Society of London B: Biological Sciences 274, 943–950.

Swaddle, J. P., Lockwood, R., 2003. Wingtip shape and flight performance in the european starling sturnus vulgaris. Ibis 145, 457–464.

Taylor, P., Lillicrap, T., Cownden, D., 2011. Inclusive fitness analysis on mathematical groups. Evolution 65 (3), 849–859.

Uecker, H., Hermisson, J., 2011. On the Fixation Process of a Beneficial Mutation in a Variable Environment. Genetics 188 (4), 915–930.

Vindenes, Y., Lee, A. M., Engen, S., Sæther, B.-E., 2009. Fixation of slightly beneficial mutations: effects of life history. Evolution 64, 1063–1075.

Wahl, L., 2011. Fixation when n and s vary: classic approaches give elegant new results. Genetics 188, 783–785.

Wahl, L. M., Dai Zhu, A., 2015. Survival probability of beneficial mutations in bacterial batch culture. Genetics 200, 309–320.

Wahl, L. M., DeHaan, C. S., 2004. Fixation probability favors increased fecundity over reduced generation time. Genetics 168, 1009–1018.

Waxman, D., 2011. A unified treatment of the probability of fixation when population size and the strength of selection change over time. Genetics 188 (4), 907–913.

Zukewich, J., Kurella, V., Doebeli, M., Hauert, C., 2013. Consolidating birth-death and death-birth processes in structured populations. PLoS One 8 (1), e54639.

